# Implicit modeling of the impact of adsorption on solid surfaces for protein mechanics and activity with a coarse-grain representation

**DOI:** 10.1101/2020.03.30.015537

**Authors:** Nicolas Bourassin, Marc Baaden, Elisabeth Lojou, Sophie Sacquin-Mora

## Abstract

Surface immobilized enzymes play a key role in numerous biotechnological applications such as biosensors, biofuel cells or biocatalytic synthesis. As a consequence, the impact of adsorption on the enzyme structure, dynamics and function needs to be understood on the molecular level as it is critical for the improvement of these technologies. With this perspective in mind, we used a theoretical approach for investigating protein local flexibility on the residue scale that couples a simplified protein representation with an elastic network and Brownian Dynamics simulations. The protein adsorption on a solid surface is implicitly modeled via additional external constraints between the residues in contact with the surface. We first performed calculations on a redox enzyme, bilirubin oxidase (BOD) from *M. verrucaria*, to study the impact of adsorption on its mechanical properties. The resulting rigidity profiles show that, in agreement with the available experimental data, the mechanical variations observed in the adsorbed BOD will depend on its orientation and its anchor residues (i.e. residues that are in contact with the functionalized surface). Additional calculations on ribonuclease A and nitroreductase shed light on how seemingly stable adsorbed enzymes can nonetheless display an important decrease in their catalytic activity resulting from a perturbation of their mechanics and internal dynamics.

## 1. Introduction

The interaction of proteins with solid surfaces is of crucial importance in the field of biomaterials^1^, since it plays a key role in numerous applications, such as tissue engineering and regenerative medicine (where we need to know the cellular response to implanted materials), the optimization of surfaces for biosensors, the development of bioactive nanoparticles, biocatalyst or bioanalytical systems.

One of the main issues that are addressed when investigating protein/surface interaction, is the matter of protein orientation. Controlled adsorption with ordered proteins is essential for devices such as biosensors, where antibodies should be immobilized with a specific orientation favoring the following antibody-antigen binding^2^, or for bioelectrocatalysis devices (biofuel cells or bioreactors for electrosynthesis), where a correct enzyme orientation is essential for direct electron transfer between the adsorbed protein and the electrode^3-4^. The adsorbed proteins’ orientation depends on many factors, such as their charge, size or shape, the support properties, or external conditions like the temperature and pH^5^. Hence these devices require a specific functionalization of the surface, in order to fine tune its charge and physico-chemical properties, that can be achieved with self assembled monolayers (SAMs) for example. For both cases, the conservation of the adsorbed proteins’ native conformation (and hence their biological function) is another key aspect that has to be taken in consideration. On the opposite end of the applications spectrum, the development of materials resisting protein adsorption has also drawn much attention, since it has practical uses, such as marine antifouling or antimicrobial coating for medical devices. In that case the challenge will be to design biomaterials where proteins will not adsorb^6-8^.

All these phenomena have now been under scrutiny for several decades, but reaching a detailed understanding of the molecular mechanisms associated with biomolecular adsorption on functionalized surfaces is far from being achieved. In particular, considerable experimental works have been conducted until now, such as atomic force microscopy, mass spectrometry and various spectroscopies^9-10^. However, the resolution of current experimental techniques is still insufficient to quantitatively determine the range of potential orientations and conformations of adsorbed proteins on the atomistic level. In that perspective, molecular simulations play an increasingly important role in revealing the mechanisms of chemical and biological processes taking place on the bio-nano interface, and designing new products. For over ten years, molecular simulation techniques have been developed to address these issues, using multiscale approaches combining all-atom and coarse-grain models^11-13^, and represent a promising tool for the biomaterials field^14-16^.

Numerous studies focus on the prediction of the adsorbed protein orientation and binding modes on the surface, which have to be controlled for example to ensure that an enzyme’s catalytic site will remain accessible after adsorption; and on the impact of adsorption on protein structure, since conformational changes are indeed likely to perturb an enzyme’s catalytic activity^17-20^. However, proteins are known for being flexible objects, and their internal dynamics play a central role for their biological function^21-22^. As a consequence, one also has to consider how surface binding will affect an enzyme’s motions to determine whether it will remain functional in its adsorbed state. In that perspective, we used a coarse-grain, elastic network representation to implicitly model the impact of surface adsorption on protein mechanics. We first performed calculations on the redox enzyme bilirubin oxidase (BOD) from the fungus *Myrothecium verrucaria*, to study the impact of adsorption on its mechanical properties. The resulting rigidity profiles show that, in agreement with the available experimental data, the mechanical variations observed in the adsorbed BOD will depend on its orientation and its anchor residues (that are in contact with the solid surface). Additional calculations on ribonuclease A and nitroreductase shed light on how seemingly stable adsorbed enzymes can nonetheless display an important decrease in their catalytic activity resulting from a perturbation of their mechanics.

## 2. Material and Methods

### Brownian Dynamics simulations

#### Rigidity profile of a protein

Coarse-grained Brownian Dynamics (BD) simulations were run using a modified version of the ProPHet (Probing Protein Heterogeneity, available online at https://bioserv.rpbs.univ-paris-diderot.fr/services/ProPHet/) program^23-25^, where additional external mechanical constraints can be applied between a set of residues defined by the user. In this approach, the protein is represented using an elastic network model (ENM). Unlike most common coarse-grained models where each residue is described by a single pseudoatom^26^, ProPHet uses a more detailed representation^27^ that involves up to 3 pseudoatoms per residue and enables different amino acids to be distinguished. Pseudoatoms closer than the cutoff parameter *Rc* = 9 Å are joined by Gaussian springs which all have identical spring constants of γstruct = 0.42 N.m^-1^ (0.6 kcal.mol^-1.^Å^-2^). The springs are taken to be relaxed for the experimentally observed conformation of the protein. Note that, following earlier studies which showed that ligands as large as a heme group actually had little influence on calculated force constants^28-29^, we chose not to include the prosthetic groups (such as metallic clusters) in the protein representation. The simulations use an implicit solvent representation via the diffusion and random displacement terms in the equation of motion^30^, and hydrodynamic interactions are included through the diffusion tensor^31^.

Mechanical properties are obtained from 200,000 BD steps at an interval of 10 fs and a temperature of 300 K. The simulations lead to deformations of roughly 1.5 Å root-mean-square deviation with respect to the protein starting conformation (which corresponds to the system’s equilibrium state). The trajectories are analyzed in terms of the fluctuations of the mean distance between each pseudoatom belonging to a given amino acid and the pseudoatoms belonging to the remaining residues of the protein. The inverse of these fluctuations yields an effective force constant *k*_*i*_ describing the ease of moving a pseudoatom with respect to the overall protein structure.

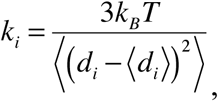

where ⟨ ⟩ denotes an average taken over the whole simulation and *d*_*i*_*=* ⟨*d*_*ij*_⟩ *j\** is the average distance from particle *i* to the other particles *j* in the protein (the sum over *j\** implies the exclusion of the pseudoatoms belonging to residue *i*). The distance between the Cα pseudoatom of residue *i* and the Cα pseudoatoms of the adjacent residues *i-1* and *i+1* are excluded since the corresponding distances are virtually constant. The force constant for each residue *k* is the average of the force constants for all its constituent pseudo atoms *i*. We will use the term *rigidity profile* to describe the ordered set of force constants for all the residues of the protein.

#### Applying external constraints on the protein

While in early work on the green fluorescent protein^32^, the external mechanical stress was modeled by applying a constant force between the Cα pseudoatoms of the residues under constraint, in this study we apply the methodology developed in ref. (33), where an external constraint is modeled by adding supplementary springs to the ENM representations. These are termed as *constraint springs*, in opposition to the structural springs resulting from the original conformation of the protein. This way we model the protein adsorption on a solid surface as a set of additional constraints that will lock the distance between protein residues which are in contact with the surface, and are termed as *anchor residues* for the rest of the manuscript. All constraint springs have the same constant γconst = 100*γstruct = 42.0 N.m^-1^ and, as for the structural springs, their equilibrium length corresponds to the original distance observed for the anchor residues in the protein experimental conformation.

#### Principal component analysis of the coarse-grained trajectories

Another useful way to investigate a protein’s internal dynamic is to look at its collective modes of motions. The BD trajectories for the proteins with and without the application of an external constraint were analyzed using principal component analysis (PCA)^34-36^ with tools from the Gromacs software package^37-39^ in order to obtain a full set of eigenvectors and eigenvalues. However, for the three proteins under study in this work, the internal dynamics are widely distributed with no dominating eigenvectors, since for all cases the 30 first eigenvectors only account for roughly 50-60% of the protein’s total variance. Subsequently, our calculations showed no clear impact of the external constraints on the proteins dynamics and this approach will not be further discussed in the manuscript.

## 3. Results and discussion

### Mechanical variations in the adsorbed bilirubin oxidase

Bilirubin oxidase is a multicopper oxidase which catalyzes the reduction of O2 to water and is considered as a promising alternative to platinum-based catalysts in enzymatic fuel cells^40-41^. Figure 1a shows the rigidity profile for the BOD from *M. verrucaria* (PDB entry 2xll) when no external constraint is applied. In agreement with earlier work on protein mechanics^29^, the most rigid residues (shown in purple in Figure 2) are located in the protein core and lie in the vicinity of the enzyme catalytic site. Residues His96, Trp132, Leu226 and Ala274 in particular are involved in the binding site of the trinuclear copper cluster (TNC) and the channels leading from the TNC to the enzyme surface^42-43^.

**Figure 1:**
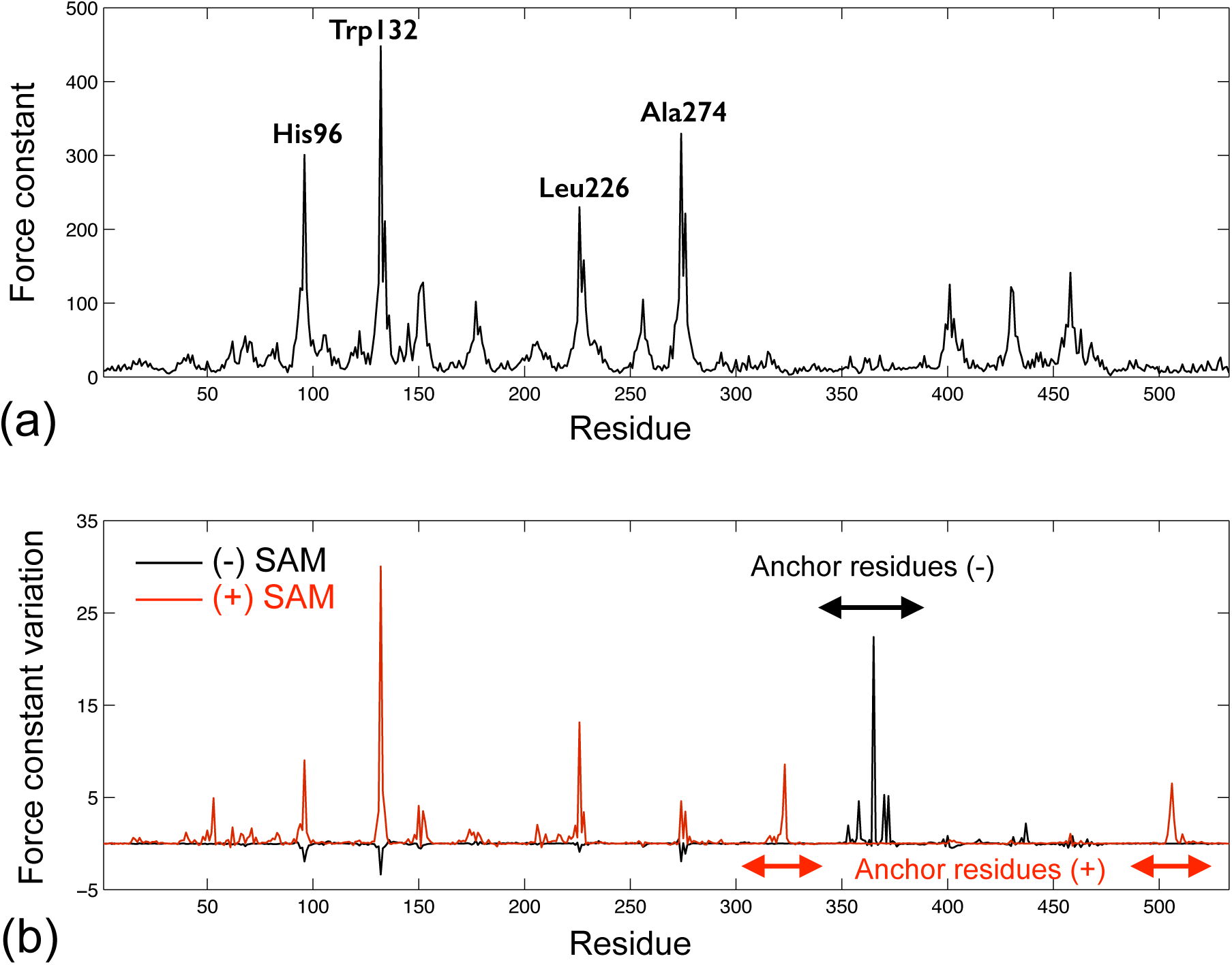
Rigidity profiles (in kcal.mol^-1^.Å^-2^) for the BOD from *M. verrucaria*. (a) Protein with no external constraint, (b) Force constant variations when applying external constraints modeling the protein adsorption on negatively charged SAMs (black line) or on positively charged SAMs (red line). The arrows highlight the position of the anchor residues (where the external constrains are applied) along the sequence.

**Figure 2:**
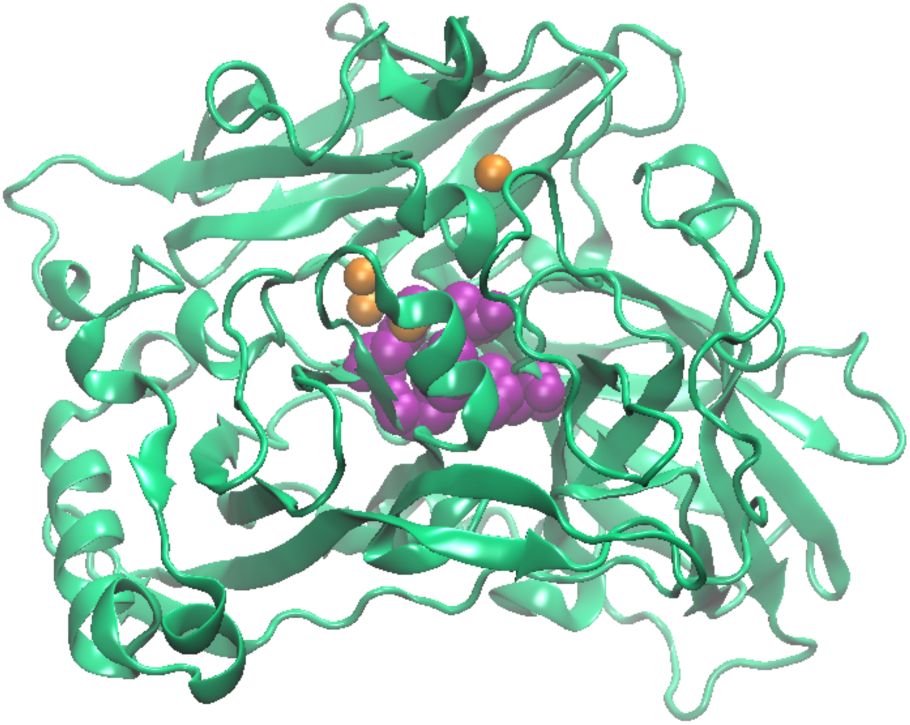
Cartoon representation for the BOD from *M. verrucaria* with the copper atoms in orange and the most rigid residues (annotated in Figure 1a) shown as purple van der Waals spheres. The trinuclear copper cluster lies next to the rigid core, while the T1 copper site is closer to the protein surface. Figure 2, 3, 4a and 5a were prepared using Visual Molecular Dynamics^60^.

To model the impact of protein adsorption on the rigidity profile, we selected as anchor residues the residues that were predicted to lie within 0.35 nm from the solid surface in the simulation work of Yang et al.44. In this study, the authors used a combination of parallel tempering Monte Carlo and all-atom Molecular Dynamics simulations to predict the orientation of BOD adsorbed on a SAM functionalized surface. Amino (-NH2) and carboxyl (-COOH) terminated SAMs were used to represent positively and negatively charged surfaces respectively, and led to different orientations of the enzyme with the following residues lying close to the surface (see Figure 3):

**Figure 3:**
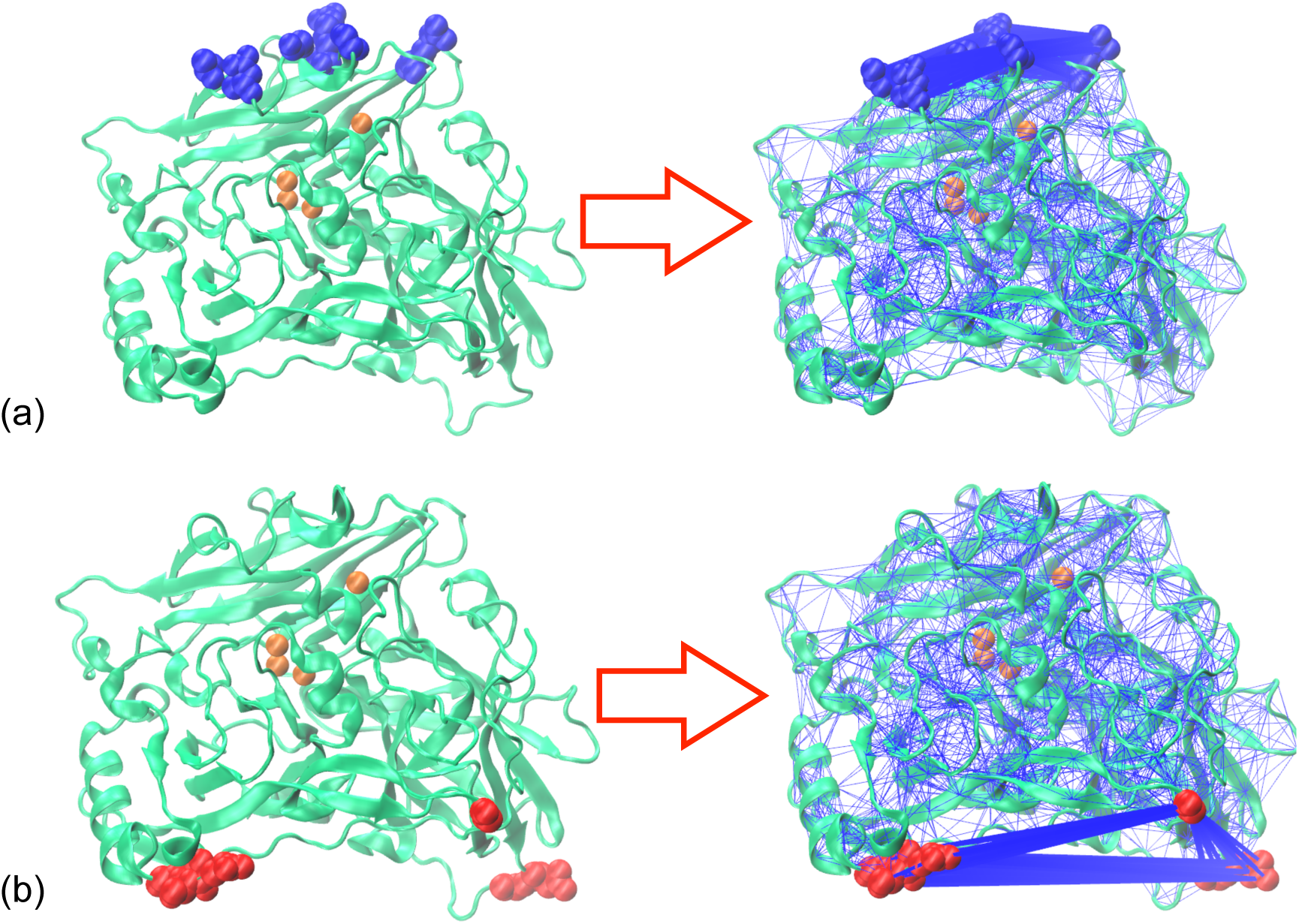
Elastic Network in our BOD CG model with the anchor residues (where additional constraint springs are added) highlighted as blue ou red van der Waals spheres. (a) Anchor residues for the negatively charged SAMs, (b) Anchor residues for the positively charged SAMs.

- Arg353, Gly358, Gly365, Asp370, Gln372 and Arg437 for the negatively charged surface.
- Asp53, Asp322, Asp323, Gln505, Ala506 and Gln507 for the positively charged surface.

The force constant variations resulting from the application of additional constraints on the anchor residues are shown on Figure 1b. For BOD oriented on the negatively charged surface (black curve in Figure 1b), the only residues that are impacted by the enzyme adsorption are the anchor residues themselves, which present an increase in their force constant, while the rest of the protein mechanics remains unchanged. In the case of a positively charged surface (red curve in Figure 1b), the protein adsorption will not only impact the anchor residues, but also residues from the central rigid core, which undergo an increase in their force constant.

In their work modeling BOD adsorption on SAMs^44^, Yang et al. focused their analyses on the interaction sites between the enzyme and the surface, and on the conformational changes undergone by the protein upon adsorption. They show that the average rmsd observed for the adsorbed BOD trajectories, 3.7 Å for COOH-SAMs and 3.0 Å for NH2-SAMs, are only slightly larger than the average rmsd observed for BOD in bulk water (2.9 Å). The observed BOD orientations on the surface are prone to favor direct electron transfer (DET) between the enzyme and negatively charged surfaces, since the T1 copper site lies closer to the electrode surface than when the enzyme is adsorbed on positively charged surfaces. However, their study concludes that the overall enzyme structure is well conserved, and does not comment on the impact of adsorption on the protein catalytic activity, including the impact on the mediated electron transfer (MET) that can be additionally or alternatively observed between BOD and the electrode. Meanwhile, Hitaishi et al. investigated the activity of the same BOD from *M. verrucaria* adsorbed on similar surfaces, i.e. gold electrodes functionalized with -COOH or -NH2 terminated SAMs, using cyclic voltammetry and spectrometric approaches (SPR, PMIRRAS and ellipsometry)^45^. In agreement with the BOD orientations predicted by Yang et al., no DET could be observed when BOD adsorption was made on the positively charged SAMs. In the case of negatively charged-SAMs, DET could be observed, with a decrease of the electron transfer rate for BOD adsorbed on 6-Mercaptohexanoic acid (6-MHA) compared to BOD on 11-mercaptoundecanoic acid (11-MUA), as a consequence of the decrease of the electron tunneling rate with the length of the alkane chain^46^. At pH = 6, the enzyme coverage was found to be similar on the negatively and positively charged surfaces, and the PMIRRAS spectra suggested that the protein general secondary structure is well conserved on both surfaces. However, the global ET rate (including both MET and DET currents) was much lower (by a factor of 4) for BOD adsorbed on the positively charged, 4-aminothiophenol (4-ATP) surface. This reduced catalytic activity for the enzyme concurs with the perturbations that we observed in the mechanical properties of the BOD adsorbed on positively charged surfaces, and which are likely to disrupt the protein function since they impact residues binding the central catalytic copper cluster. While leaving the global structure intact, the external constraints applied on anchor residues upon adsorption increase the rigidity of functional residues that lie between 20 and 25 Å away from them. Note that this long range effect of mechanical perturbations has already been observed in earlier studies on bacterial reaction centers^47-48^.

### Adsorbed ribonuclease A

Ribonuclease A (RNase A) is a small (120 residues) enzyme that catalyzes the breakdown of the phosphodiester backbone in RNA. Its immobilization on solid surfaces is of interest for applications such as the purification of biopharmaceutical plasmid DNA^49^, and it is also currently investigated as a potential anti tumoral agent that could be coupled to material surfaces in various drug delivery platforms^50^. In their experimental work^51^, Wei et al. investigated the enzymatic activity of RNase A adsorbed on three different surfaces, namely silica glass, high density polyethylene (HDPE) and poly(methyl-methacrylate) (PMMA). Adsorption caused RNase A to undergo a substantial degree of unfolding and to lose about 60% of its native-state catalytic activity, independently of the material it was adsorbed on. In their theoretical study using MD simulations, Liu et al. investigated the behavior of RNase A adsorbed on SAM-functionalized surfaces with opposite charges (thus leading to opposite enzyme orientations)^52^. In both cases, they observed that the enzyme native conformation is preserved. Here we modeled the impact of adsorption on RNase A mechanical properties for four different materials, using as starting point the high resolution crystal structure with the PDB id 7rsa, and as anchor residues those mentioned in the aforementioned studies (see Figure 4a):

**Figure 4:**
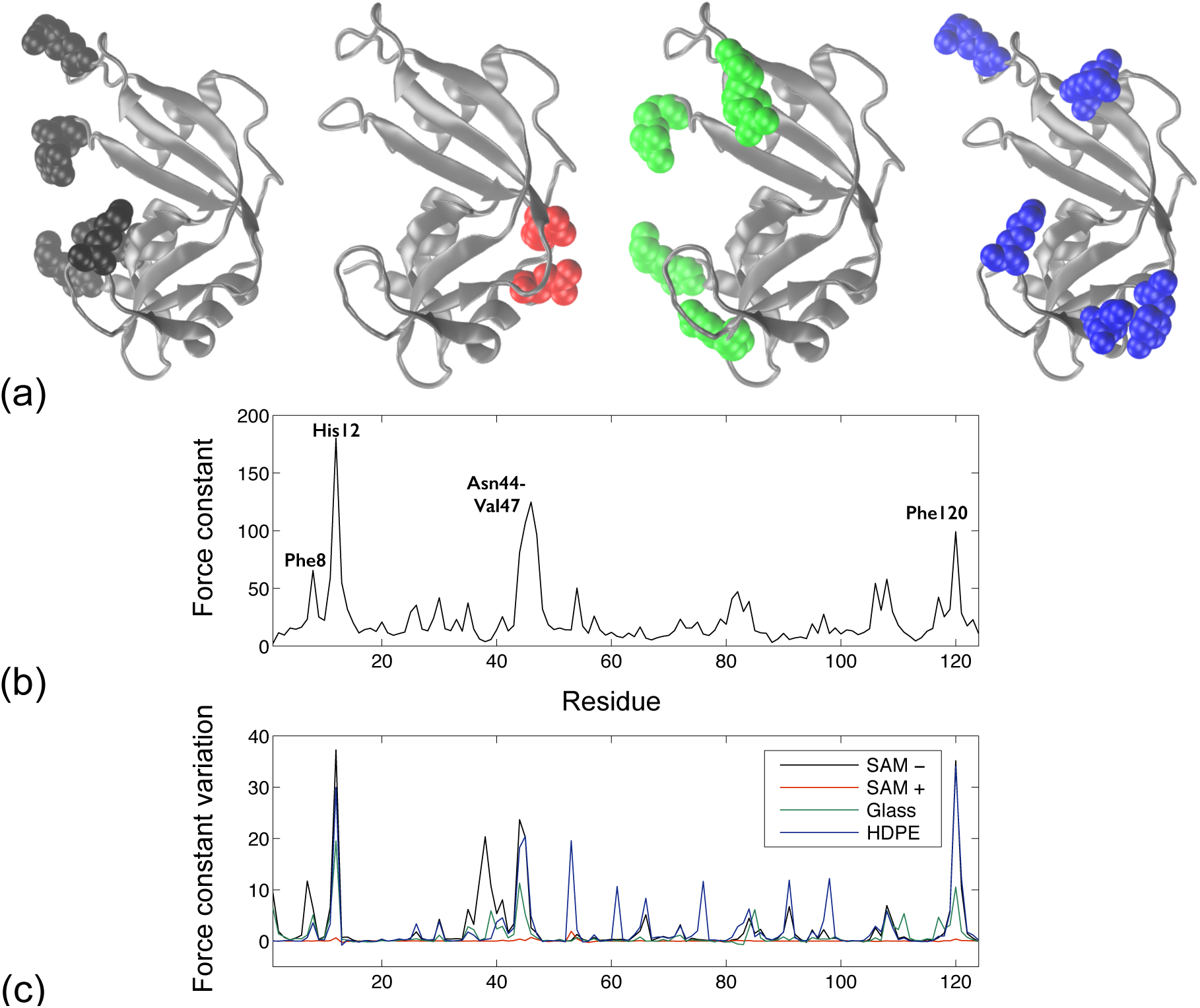
(a) Cartoon representation for RNase A (PDB id 7rsa) with the anchor residues shown as van der Waals spheres for different surfaces: Negatively charged SAMs (black), positively charged SAMs (red), glass (green) and HDPE (blue). (b) Rigidity profile (in kcal.mol^-1^.Å^-2^) for the RNase A from *B. taurus* with no external constraints. (c) Force constant variation when applying external constraints modeling the protein adsorption with the anchor residues shown in panel (a) and with the same color code.

- Negatively charged SAMs (SAM-), Lys1, Lys7, Lys37, Asp38, Arg39, Lys66, Lys91
- Positively charged SAMs (SAM+), Glu49, Asp53
- Glass, Lys1, Arg39, Arg85, Glu111 (leading to the same enzyme orientation as the SAM-surface)
- HDPE (and also PMMA), Asp53, Lys61, Lys66, Tyr76, Lys91, Lys98

For the SAM+ orientation, where there are only two anchor residues which lie very close to each other, the adsorption has little to no impact on the enzyme rigidity profile (see the red line on Figure 4c). For all the other materials, adsorption induces a strong increase in the force constant of a limited set of residues. This set includes the anchor residues, as could be expected, but also residues from the enzyme catalytic site (His12 and Lys41)^53^, and residues forming the protein folding nucleus (Phe8, Phe46, Val47 and Phe120)^54^. Disrupting the dynamic properties of these to groups of residues is indeed likely to impact the catalytic activity of RNase A.

### Adsorbed nitroreductase

Nitroreductase (NfsB) is a FMN-dependent homodimeric enzyme which catalyzes the reduction of a wide range of substrates containing nitro-groups. It is well characterized both structurally and functionally, and often serves as a model system to study how surface interactions will affect enzyme stability and activity^55-58^. Recently, Zou et al. investigated the impact of two-point surface attachment on protein function by producing NfsB mutants with neighboring tethering sites and using spectroscopic approaches to deduce their orientation and stability^58^. Interestingly, two mutants, with the anchor residue positions (111, 423) and (247, 423), display better structural stability than any single-site immobilized enzyme. However, these mutants are also less active than the single-site ones thus suggesting that there is a trade-off between activity and stability. These experimental results concur with our coarse grain simulations on NfsB from *E. coli* (PDB: 1ds7), where both two-sites constraints induce changes of the residues force constants in the protein rigid core, see Figure 5. On Figure 5c, we can see how applying an external constraint between anchor positions (111, 423) or (247, 423) disturbs the original periodicity observed in the rigidity profile of the unconstrained enzyme in Figure 5b (which reflects the protein homodimeric structure). Unlike the previous cases for BOD and RNase A where the additional constraints modeling protein adsorption would only induce an increase in the protein rigidity, the constraint between the anchor residues in NfsB leads to a mixed mechanical response, with residues from the central rigid core undergoing either a decrease or an increase of their force constant. The disruption of the initial pattern observed in Figure 5b may be related to the decrease in activity observed experimentally in ref. (58).

**Figure 5:**
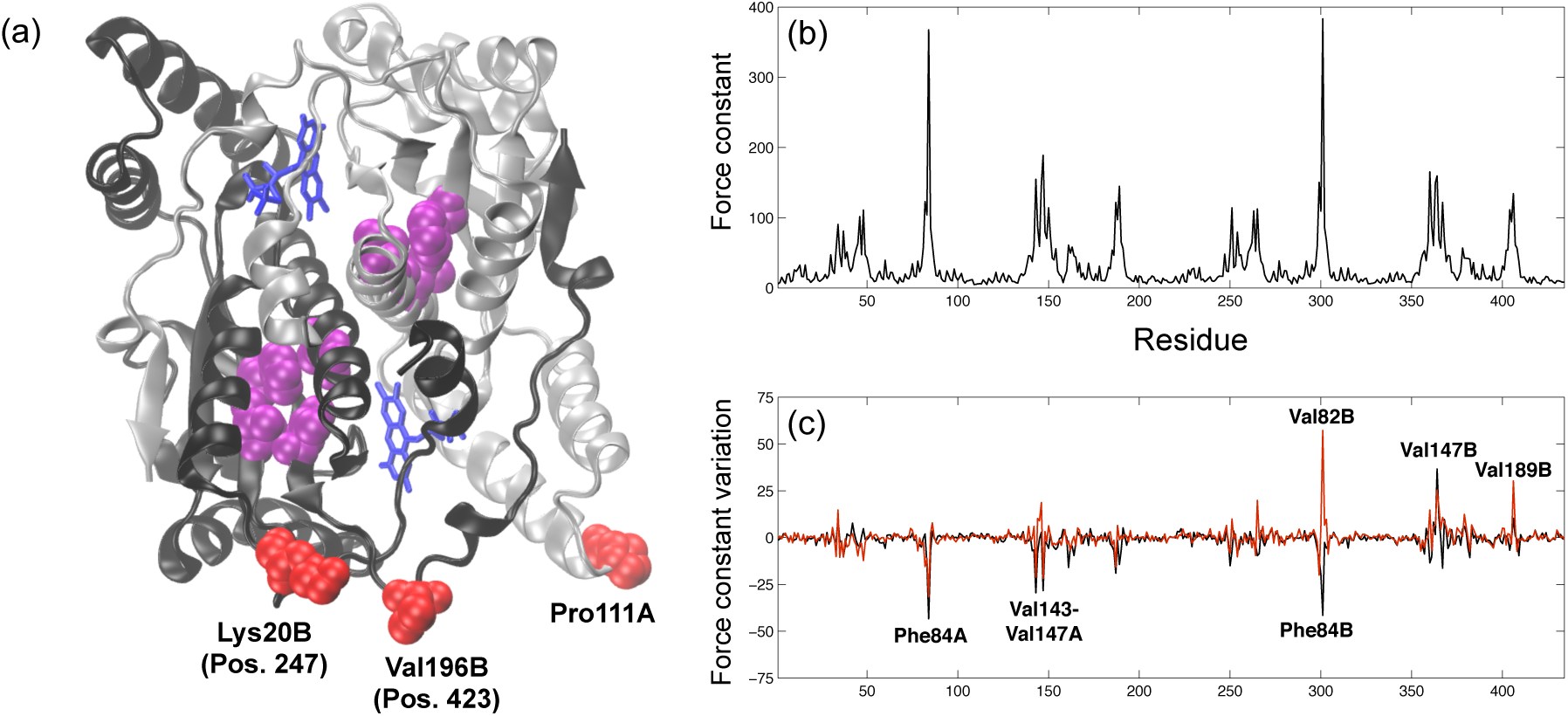
(a) Cartoon representation for the homodimeric NfsB (PDB id 1ds7) with FMN cofactors in blue. The anchor residues are shown as red van der Waals spheres, and the residues undergoing the largest force constant variations upon surface adsorption are in purple. (b) Rigidity profile (in kcal.mol^-1^.Å^-2^) for the homodimeric NfsB with no external constraints. (c) Force constant variation when applying and external constraint between anchors positions (423,111), black line, or (423,247), red line.

## 4. Conclusion

Enzymes immobilized on solid surfaces appear in numerous devices that are currently developed for a wide range of applications, such as biocatalysis, biosensing or healthcare. However, the efficient integration of enzymes in nanomaterials is a complicated issue since it requires the simultaneous preservation of the enzyme structure, dynamics and catalytic site accessibility, a goal that remains difficult to achieve. In this study, we use a modified version of the ProPHet program, which combines a coarse-grain protein representation and Brownian Dynamics simulations, to investigate the impact of surface immobilization on protein mechanics. Protein adsorption is modeled in an implicit manner by adding supplementary constraints between the anchor residues, i. e. residues in contact with the solid surface, to the global elastic network. This permits to observe the mechanical impact of adsorption even though the protein structure remains intact during the process. The first calculations were done on a bilirubin oxidase, an enzyme involved in the development of biofuel cells, and they show that the mechanical perturbations resulting from these additional constraints depend on the enzyme’s orientation on the surface and the resulting set of anchor residues. Furthermore, the rigidity increase observed in the protein following its adsorption on a positively charged surface not only concerns the anchor residues (as could be expected), but it can also be a long range effect, and impact the force constants of residues located in the enzyme active site. This, in turn, is likely to be detrimental for the catalytic function, and our results concur with the experimental observations made by Hitaishi et al, who compared the electron transfer rates and conformation of BOD adsorbed on positively and negatively charged surfaces.

Further calculations on the enzymes RNase A and nitroreductase, for which experimental data regarding the impact of adsorption on function is available, lead to similar conclusions; i. e. the experimentally observed activity loss upon surface binding can be associated with mechanical perturbations of the catalytic residues.

Globally, our results highlight the fact that the structural integrity of surface immobilized enzymes is no guarantee that the enzyme will remain functional, and that the systems’ internal dynamics should be specifically investigated before drawing any conclusions regarding the conservation of the catalytic activity. The mechanical perturbations obtained for RNase A grafted on various surfaces (and with various orientations) illustrate the stability-activity tradeoff that was recently observed by Weltz et al.^59^ using single-molecule FRET imaging. Altogether our modeling approach provides a simple tool to help understand how protein adsorption might impact their dynamics and function, especially when one needs to achieve a delicate balance between an enzyme stability and its catalytic activity.

## Acknowledgments

This work was supported by the ANR (ENZYMOR-ANR-16-CE05-0024) and by the “Initiative d’Excellence” program from the French State (Grant “DYNAMO”, ANR-11-LABX-0011-01).

---

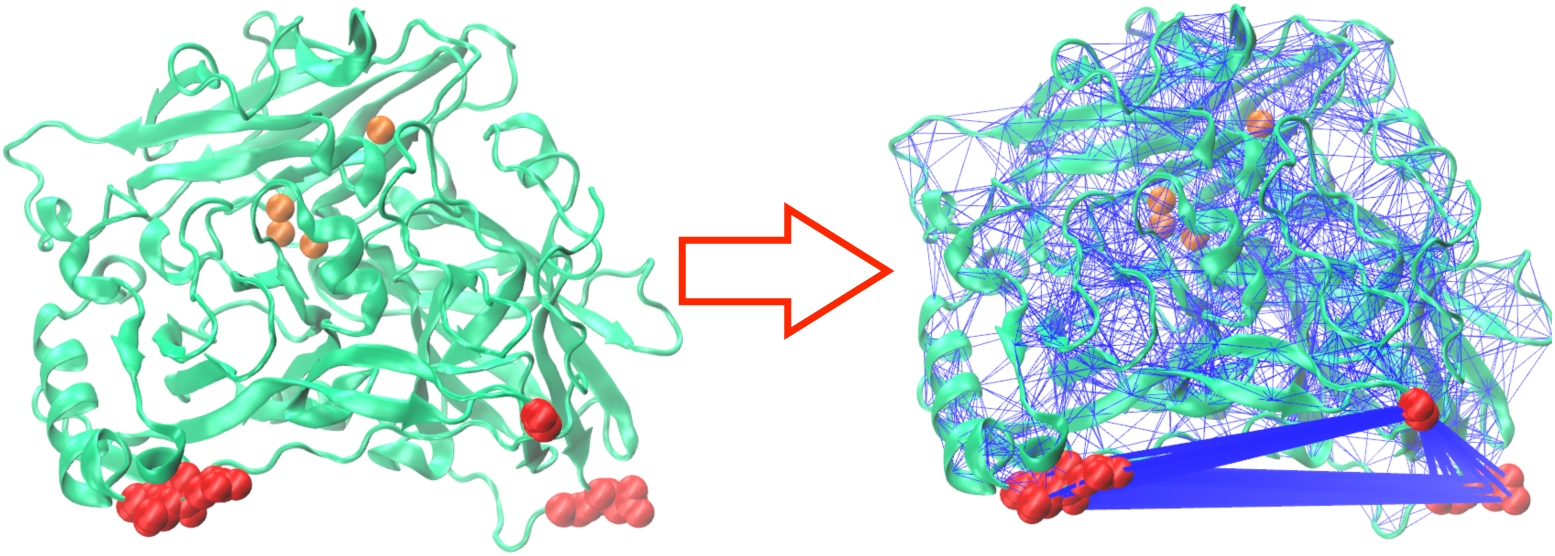

**TOC image**

